# PPAR-γ activation promotes xenogenic bioroot regeneration by attenuating the xenograft induced-oxidative stress

**DOI:** 10.1101/2022.03.29.486266

**Authors:** Tingting Lan, Fei Bi, Yuchan Xu, Xiaoli Yin, Jie Chen, Xue Han, Weihua Guo

**Author notes:** Corresponding author at: Department of Pediatric Dentistry, West China School of Stomatology, Sichuan University, No. 14, 3rd Sec., Ren Min Nan Road, Chengdu 610041, P.R. China, E-mail address (W. Guo).

## Abstract

**Objective:** xenogenic organ transplantation has been considered the most promising strategy in providing possible subtitutes with physiological function of the failing organs as well as solving the problem of insufficient donor sources. However, the xenograft, suffered from immune rejection and ischemia-reperfusion injury (IRI), causes massive ROS expression and the subsequent cell apoptosis, leading to the xenograft failure. Our previous study found a positive role of PPAR-γ in anti-inflammation through its immunomodulation effects, which inspires us to apply PPAR-γ agonist rosiglitazone (RSG) to address survival issue of xenograft with the potential to eliminate the excessive ROS.

**Methods:** **X**enogenic bioroot was constructed by wrapping the dental follicle cells (DFC) with porcine extracellular matrix (pECM). The hydrogen peroxide (H_2_O_2_)-induced DFC was pretreated with RSG to observe its protection on the damaged biological function. Immunoflourescence staining and transmission electron microscope were used to detect the intracellular ROS level. SD rat orthotopic transplantation model and SOD1 knockout mice subcutaneous transplantation model were applied to explore the regenerative outcome of the xenograft.

**Results:** RSG pretreatment significantly reduced the adverse effects of H_2_O_2_ on DFC with decreased intracellular ROS expression and alleviated mitochondrial damage. In vivo results confirmed RSG administration substantially enhanced the host’s antioxidant capacity with reduced osteoclasts formation and increased periodontal ligament like tissue regeneration efficiency, maximumly maintaining the xenograft function.

**Conclusions:** RSG preconditioning could preserve the biological properties of the transplanted stem cells under OS microenvironment and promote organ regeneration by attenuating the inflammatory reaction and OS injury.

## Introduction

In future, porcine derived ECM will be the most widely used bio-scaffold for ECM-based bioengineering transplantation to replace the failing organs and solve the problem of the limited donor source (***Chen et al., 2020; Lan et al., 2021***). However, the application of xECM-based organ transplantation faces great challenges for the transplanted xenografts are often susceptible to redox imbalance and immune rejection due to their heterologous antigens which greatly aggravate the nonspecific inflammatory response and subsequent overproduction of ROS. (***Choi et al., 2015; Lan et al., 2021***).

Appropriate ROS concentration is vitally important in a variety of pathophysiological conditions by maintaining the microenvironment homeostasis through eliminating inflammatory mediums when injury occurs. Excessive intracellular ROS will amplify the inflammatory response (***Maccarrone et al., 2009; Lei et al., 2015***), which will induce genome instability with intracellular DNA damages through activating nuclear factor-kappa B (NF-κB) pathway in macrophages, promoting the macrophages polarizing to M1 phenotype (***Seebach et al., 2019; Vallée et al., 2018; Nauta et al., 2014***), directly causing the transplanted stem cells’ death and multipotency impairment (***Maccarrone et al., 2009; Lei et al., 2015; Ott et al., 2018; Vono et al., 2018***). Moreover, abundant M1 polarized macrophages will increase the secretion of inflammatory factors, such as tumour necrosis factor (TNF)-α and interleukin (IL)-1 β, which could further recruit the neutrophils around the transplantation site and thus, aggravate inflammatory reaction (***Barshes et al., 2005***). In addition to the excessive ROS production, the decreased intracellular antioxidant capacity with down-regulated expression of superoxide dismutase (SOD), glutathione peroxidase (GPx) and glutathione (GSH) also contributes to redox imbalance in the xenograft transplantation process (***Cappelli et al., 2005***), making the tissue more vulnerable to OS and inflammatory reaction. The aggravated inflammation in the microenvironment could cause great damage to the cells, contributing to the xenograft failure. For example, when conducting the pancreatic islet transplantation, it is usually accompanied by up-regulated expression of ROS related to the nonspecific inflammatory response (***Eltzschig et al., 2011; Alva et al., 2018; Zhang et al., 2021***), immune rejection and IRI with persistent redox imbalance (***Yamada et al., 2013***), which would cause the islet dysfunction and unable to regulate blood glucose in type 1 diabetes mellites treatment.

Given the importance of OS and inflammatory response in the survival of the transplanted organ, it is of great importance to suppress ROS expression to alleviate the inflammation after the procedure of transplantation.It has been reported that when the excessive ROS is scavenged (***Barshes et al., 2005***), the regeneration process of the destroyed tissue in islet graft can be restarted to some extent. An emphasis on ROS control under the situation of promoting renal transplantation survival has been widly proposed during last 20 years since the most serious complications in kidney transplantation often result from OS (***Tejchman et al., 2021; Farina et al., 2019; Yao et al., 2020***). Similarly, during ECM based bioroots transplantation process, application of antioxidant N-Acetyl-L-cysteine (NAC) has obviously improved the periodontal ligaments (PDLs) regeneration efficiency by preserving the biological function of the transplanted stem cells, verifying the importance of antioxidation in bioroots transplantation (***Zhang et al., 2021***). In order to acquire an favorable outcome of xenogenic bioroots transplantation, not only controlling the overproduction ROS is essential, but enhancing the host’s ability to defend against immune rejection is also an imperative requirement. Therefore, co-transplantation of xenogenic bioroots with a drug that could mediate the immune evasion as well as alleviate the OS damage is a promising strategy for xenogenic organ survival.

Peroxisome proliferator-activated receptor γ (PPARγ) receptor is an important signal molecule in regulating various physiological and pathological process like inflammation, immune reaction and cell metabolism (***Evans et al., 2014; Lu et al., 2011; Stienstra et al., 2008; Charo, 2007***). Recently, PPARγ agonist rosiglitazone (RSG), has been proved to be served as an effective immunosuppressive agent, which shows superiority on modulating immune microenvironment by promoting M2 macrophage polarization during acute immune rejection of the perioperative period of xenogenic organ transplantation (***Lan et al., 2021***). Moreover, RSG also shows strengths in inflammation controlling and neuron preservating during renal transplantation (***Elshazly et al., 2019***), making it a good candidate for microenvironment regulation of xenobiotic roots.

Our previous studies verified the efficiency of RSG on immunomodulation with the potential of alleviating the early-stage immune rejection (***Lan et al., 2021***). Therefore, in this study, we focused on exploring the protective effect of RSG on the intracellular environmental homeostasis of the transplanted stem cells, referring as DFCs in the xenogenic bioroots system under oxidative stress. We constructed an in vitro OS model with H_2_O_2._ SD rat orthotopic transplantation model was established to explore whether the antioxidant effect of RSG on the transplantation microenvironment could promote the final regeneration results of the xenogenic bioroots. For further verifying the role of RSG in tissue remodeling process under OS, xenogenic bioroot was cultured in subcutaneous areas of SOD1 knockout (SOD1^-/-^) mice. We discovered a positive effect of RSG on suppressing ROS expression, resulting in a promotion on the implanted organ survival. This approach demonstrates a potential for xenogenic tissue or organ regeneration with PPARγ.

## Results

### Special biological properties of pTDM and rDFC were conducive to construct xenogenic bioroots

Compared to pNDM, pTDM showed a porous and lax structure, through which proteins could be easily released (***Figure 1A***). Large amounts of protein were continuously released into the culture supernatant in the first five days of pTDM culture and its concentration decreased significantly along incubation time (***Figure 1B***). Additionally, a significant promotion effect on the cell’s viability by pTDM protein, either in higher or lower concentration, was detected (***Figure 1C***). In order to further confirm the kinds of proteins released from pTDM, ELISA assay was conducted. It showed a sustained release mode with a relative high concentration of the osteogenic or odontogenic differentiation-related proteins like Biglycan, bone glaprotein (BGP / OCN), TGF-β 1, dentin matrix protein 1 (DMP-1) and dentin sialoph-osphoprotein (DSPP) within 4 weeks (***Figure 1D***). Next, characteristics of rDFC were explored. A large number of spindle shaped cells migrated from the harvested tissue could be seen (Figure 1E and 1I). The morphology of rDFC shifted towards fuzziness when extracellular matrix membrane formed (***Figure 1E***). IF staining with positive vimentin and negative CK14 confirmed that the harvested cells were dental follicle mesenchymal stem cells rather than epithelial cells (***Figure 1F***). At last, multiple differentiation potential of rDFC was determined with a large number of mineralized nodules formation by Alizarin red staining (ARS) after culturing in the osteogenic induction medium for 21 days and positively stained with β III tubulin by IF staining after culturing in the neurogenic induction medium for 2 hours (***Figure 1G and 1H***). Above results suggested that rDFC wrapped with pTDM was a feasible strategy for constructing the xenogenic bioengineered roots and highlighted that pTDM’s sustained biological proteins releasing had made it an ideal bio-scaffold for inducing the odontogenic differentiation of rDFC.

**Figure 1.**
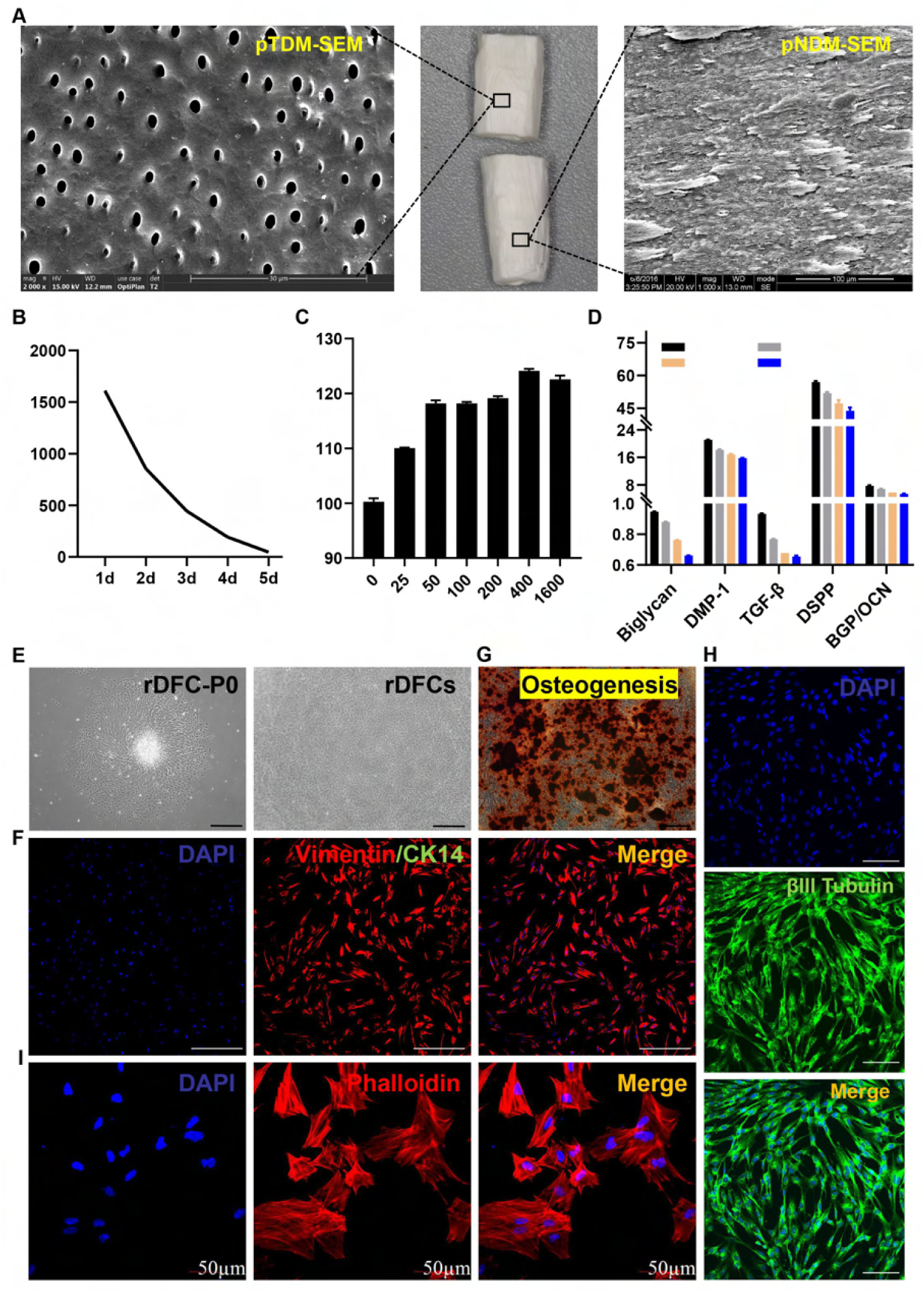
Special characteristics of pTDM and rDFC were conducive to construct xenogenic bioroots. A, SEM image of pTDM and pNDM. B, concentrations of the protein released from pTDM at different culture time points. C, effect of the different concentrations of pTDM protein on rDFC’s viability. **P < 0.01; ***P < 0.001. D, expressions of Biglycan, BGP/OCN, TGF-β1, DMP-1 and DSPP in the pTDM extracts tested by ELISA kits. E, rDFC morphology observation with light microscope. Scale bar = 500μm. F, IF staining with Vimentin and CK14 to determine the source of cells. Scale bar = 200μm. G, ARS staining of rDFC cultured in the osteogenic induction medium for 14 days. Scale bar = 500μm. H, IF staining with βIII-Tubulin of rDFC cultured in the neurogenic induction medium for 2 hours. Scale bar = 100μm. I, IF staining with phalloidin of rDFC to determine cell morphology under laser confocal scanning microscope.

### RSG reversed the biological characteristics of H_2_O_2_-induced damaged cells in the xenogenic bioroot system with PPAR-γ receptor activation

To determine the optimum inducing concentrations, CCK-8 test was conducted after rDFC was cultured with different concentrations of H_2_O_2_ or RSG. As shown in ***Figure 2A***, over 25uM H_2_O_2_ could directly induce large amounts of cells death and furthermore, over 10uM RSG could lead to obvious decreased cell survival as shown in ***Figure 2B***. Therefore, 25uM H_2_O_2_ was selected as working concentration to induce cellular oxidative stress while 10uM RSG was used to explore the effect of RSG on the H_2_O_2_-induced damaged cells. It showed that cell survival increased slightly from 83% to 94% with RSG pre-treated the H_2_O_2_ induced cells (***Figure 2C***). In addition to the promotion on the cells viability, RSG also showed an improvement on cell migration. As shown in ***Figure 2D***, compared to the H_2_O_2_-induced cells, RSG pretreated cells showed an enhancement on the cell’s migration at 24 h and 48 h, indicating the protective role of RSG on the migration ability of cells exposed to the oxidative stress (***Figure 2D***). In order to further explore the effect of RSG on the osteogenic/odontogenic differentiation potential of rDFC treated with H_2_O_2_, ARS staining and RT-PCR experiment were carried out. Results of RT-PCR showed that compared to the control group (rDFC), the osteogenic/odontogenic differentiation ability of cells treated with H_2_O_2_ was significantly impaired, accompanied by down-regulated expression of ALP and COL-1. However, administration with RSG could partially reverse the inhibitory effect of H_2_O_2_ on osteogenic differentiation of cells, preserving its biological function close to the normal cells (***Figure 2E***). Similarly, the number of mineralized nodules stained by ARS in cells stimulated by H_2_O_2_ significantly decreased after culturing in the osteogenic inducing medium for 21 days while pretreatment with 10 uM RSG could partially block this effect (***Figure 2F***). These results suggested that RSG could protect the biological characteristics of H_2_O_2_-induced damaged cells in the xenogenic bioroot system with PPAR-γ receptor activation.

**Figure 2.**
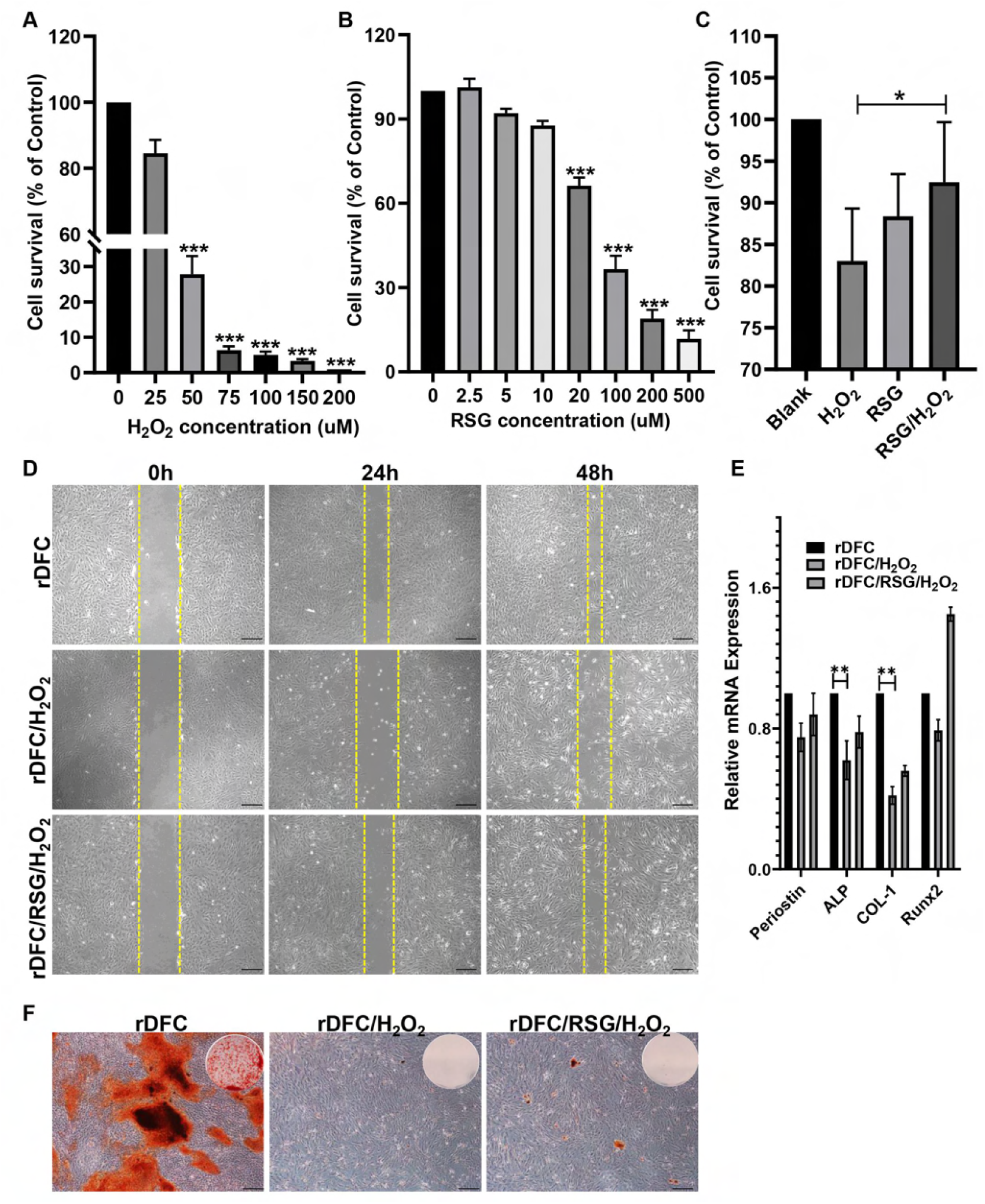
RSG protected the biological characteristics of H_2_O_2_-induced damaged cells in the xenogenic bioroot system with PPAR-γ receptor activation. A and B, effects of different concentrations of H_2_O_2_ and RSG on cells viability. *p < 0.05; **p < 0.01; ***p < 0.001. C, effect of 10uM RSG on the proliferation of 25 uM H_2_O_2_ induced-rDFC. D, migration efficiency of the rDFC under different stimulation at 0 h, 24 h and 48h. Scale bar = 500μm. E, relative mRNA expression of Periostin, ALP, COL-1 and Runx2 at 7 days. **p < 0.01. F, effect of RSG on the osteogenic differentiation potential of the H_2_O_2_-induced rDFC as determined by the amount of the mineralized nodules through ARS staining. Scale bar = 500μm.

### RSG alleviated the intracellular oxidative stress status in vitro

For exploring the intracellular ROS level of rDFC, DCFH-DA, JC-1 staining and TEM observation were conducted. In DCFH-DA staining, there was an obvious elevated ROS expression of rDFC induced by H_2_O_2_ while an significant decreasing ROS expression of rDFC pretreated with10uM RSG, judged by fluorescence intensity (***Figure 3A***). Consistent with this result, mitochondrial membrane potential detected by JC-1 staining also showed a relatively low mitochondrial membrane potential of rDFC induced by H_2_O_2_ while a higher level of mitochondrial membrane potential of rDFC with RSG pretreatment, judged by red fluorescence intensity, confirming the positive role of RSG in controlling ROS expression (***Figure 3B***). TEM observation showed highly swollen mitochondria with mitochondrial membrane rupture, mitochondrial ridge continuity broken, even disappear and peripheral condensation of chromatin of the rDFC with H_2_O_2_ treated. When pretreated with RSG, the ultrastructure of mitochondria of rDFC tended to be normal, verifying the protective role of RSG on mitochondria exposed to H_2_O_2_-induced OS (***Figure 3C***). For evaluating the antioxidant capacity of cells, total antioxidant capacity of rDFC stimulated with different chemical regents were determined. As shown in Figure 3D, when treated with H_2_O_2_, the cells antioxidant capacity decreased significantly from 181uM/g to 135uM/g while it elevated to 216 uM/g with RSG pretreating the H_2_O_2_-induced cells (***Figure 3D***). To verify the intracellular inflammatory level, IL-4 concentration was determined by elisa test. As shown in Figure 3E, compared to the control group with 0.607pg/ml, IL-4 expression of the H_2_O_2_-induced cells decreased obviously while RSG pretreatment could promote its concentration to 0.321pg/ml (***Figure 3E***).

**Figure 3.**
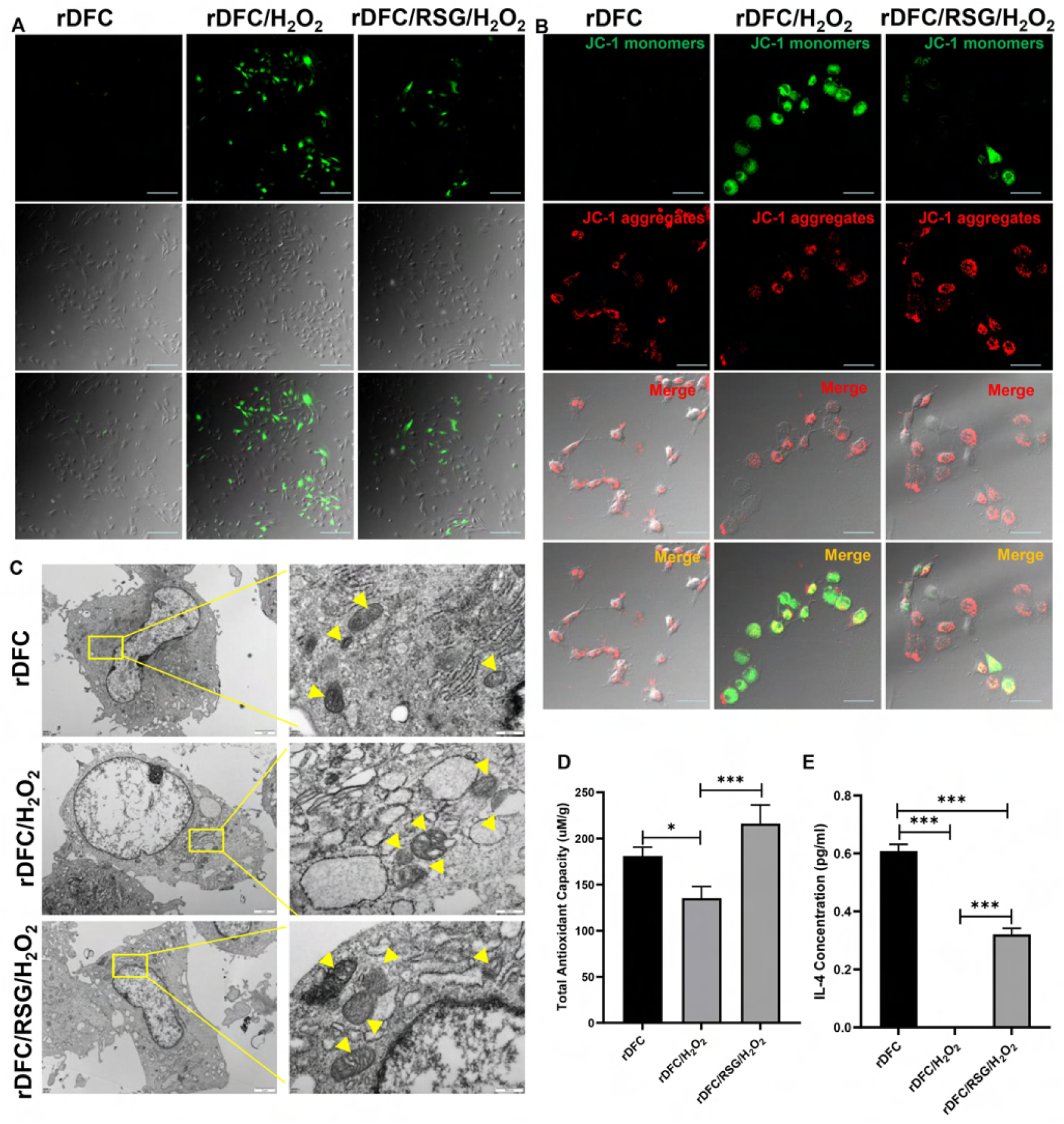
RSG alleviated the intracellular oxidative stress status in vitro. A, intracellular oxidative stress detected by DCFH-DA. Scale bar = 100μm. B, intracellular oxidative stress detected by JC-1. Scale bar = 50μm. C, ultrastructure of mitochondria observed by TEM. yellow arrow: mitochondria. D, total antioxidant capacity of rDFC stimulated with different chemical regents. *p < 0.05; ***p < 0.001. E, IL-4 concentration in the cell supernatant detected by elisa test. ***p < 0.001.

### RSG suppressed xenograft induced-oxidative stress

To explore whether RSG could be effective for improving the microenvironment of xenografts by controlling ROS level, orthotopic implant model of SD rats was built by transplanting the xenogenic bioengineered tooth organs into the alveolar fossa (***Figure 4A***). Compared to pTDM-rDFCs group, concentration of 8-OHdG (8-hydroxydeoxyguanosine) in pTDM-rDFCs/RSG group significantly decreased from 29.68 ng/mL to 22.75 ng/mL, although the level of 3-NT (3-nitrotyrosine) had no significant difference between the two groups (***Figure 4B, 4C***). Consistent with this, HNEJ-2 (4-hydroxynonenal, a product of lipid peroxidation) expression around the transplanted xenogenic bio-roots at 1 week post implantation significantly down-regulated in pTDM-rDFCs/RSG group. However, at 1 month post surgery, both of the groups with or without RSG treatment exhibited a lower HNEJ-2 level (***Figure 4D***), indicating the effective role of RSG in suppressing the OS injury in the early stage post transplantation.

**Figure 4.**
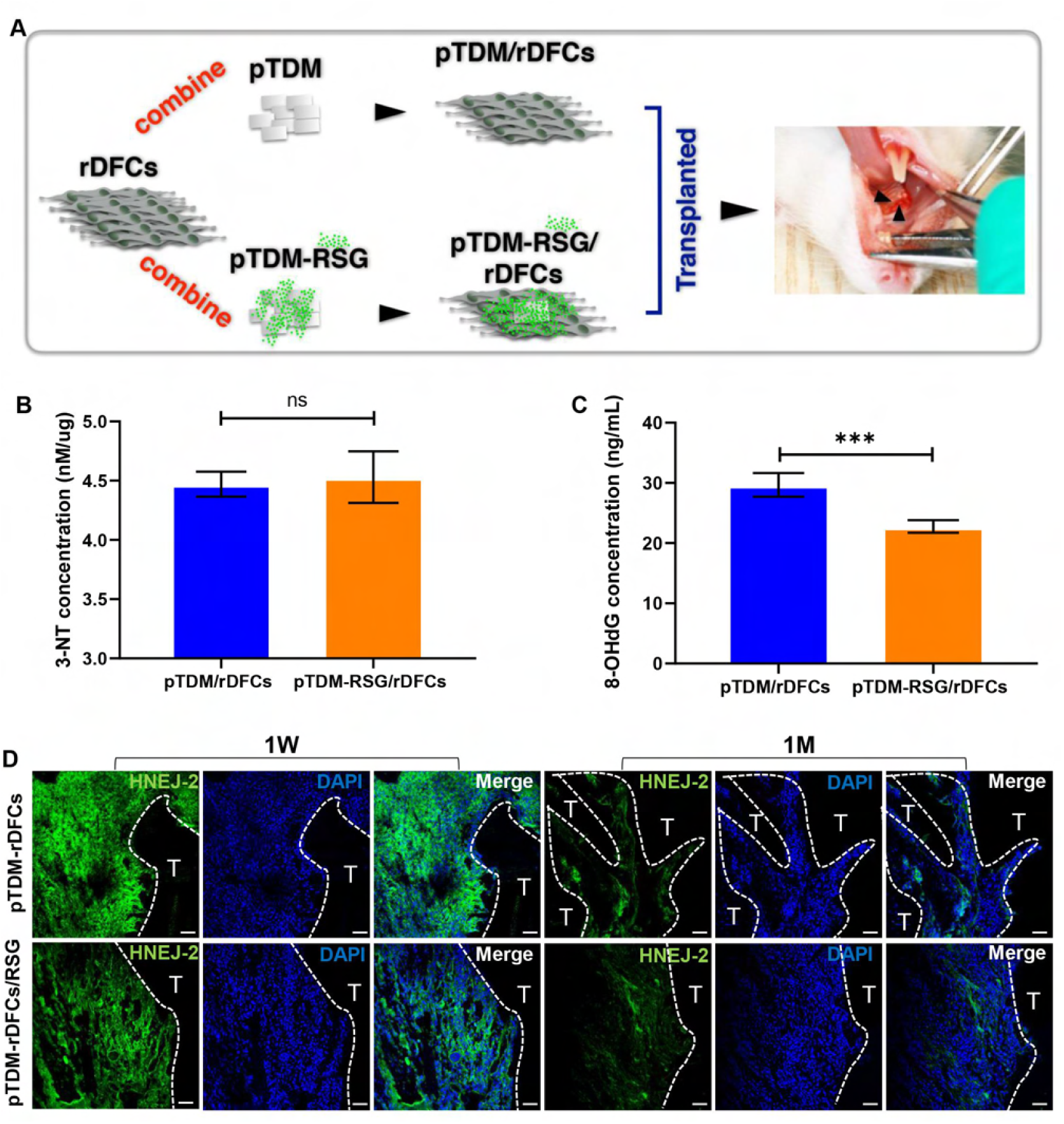
RSG suppressed xenograft induced-oxidative stress. A, schematic illustration of orthotopic transplantation of SD rats. B and C, Expressions of 3-NT and 8-OHdG around the xenogenic bio-roots in vitro tested by Elisa. ns: no significant difference; ***P < 0.001. D, IF staining of HNEJ-2 (green) around pTDM at 1 week and 1 month for determining ROS level. T: pTDM. Scale bar = 100μm.

### RSG inhibited osteoclasts differentiation while promoted PDL-like tissue regeneration in xenogeneic bioroots remodeling process by inhibiting inflammatory reaction

For detecting the effect of RSG on inflammation controlling and the regeneration results, IF staining with TNF-α, histological staining and trap staining of the samples were performed. It showed a significant decreased expression of TNF-αwith RSG treatment at 1 week post implantation, which accorded with the results in HNEJ-2 staining (***Figure 5A***). At 2 months after implantation, it showed an active bone remodeling response in xenogenic bioroots system, including obvious bone adhesion and bone absorption, with large amounts of osteoclasts differentiation around pTDM. Consequently, no obvious collagen fibers regeneration was observed. However, with RSG treatment, lots of collagen fibers, arranged perpendicular to pTDM surface similar to natural PDL, were regenerated without obvious bone adhesion or pTDM resorption. Besides, less osteoclasts were detected around pTDM (***Figure 5B***). These results suggested that RSG could significantly inhibit osteoclasts formation but promote PDL-like tissue differentiation in xenogenic bioroots.

**Figure 5.**
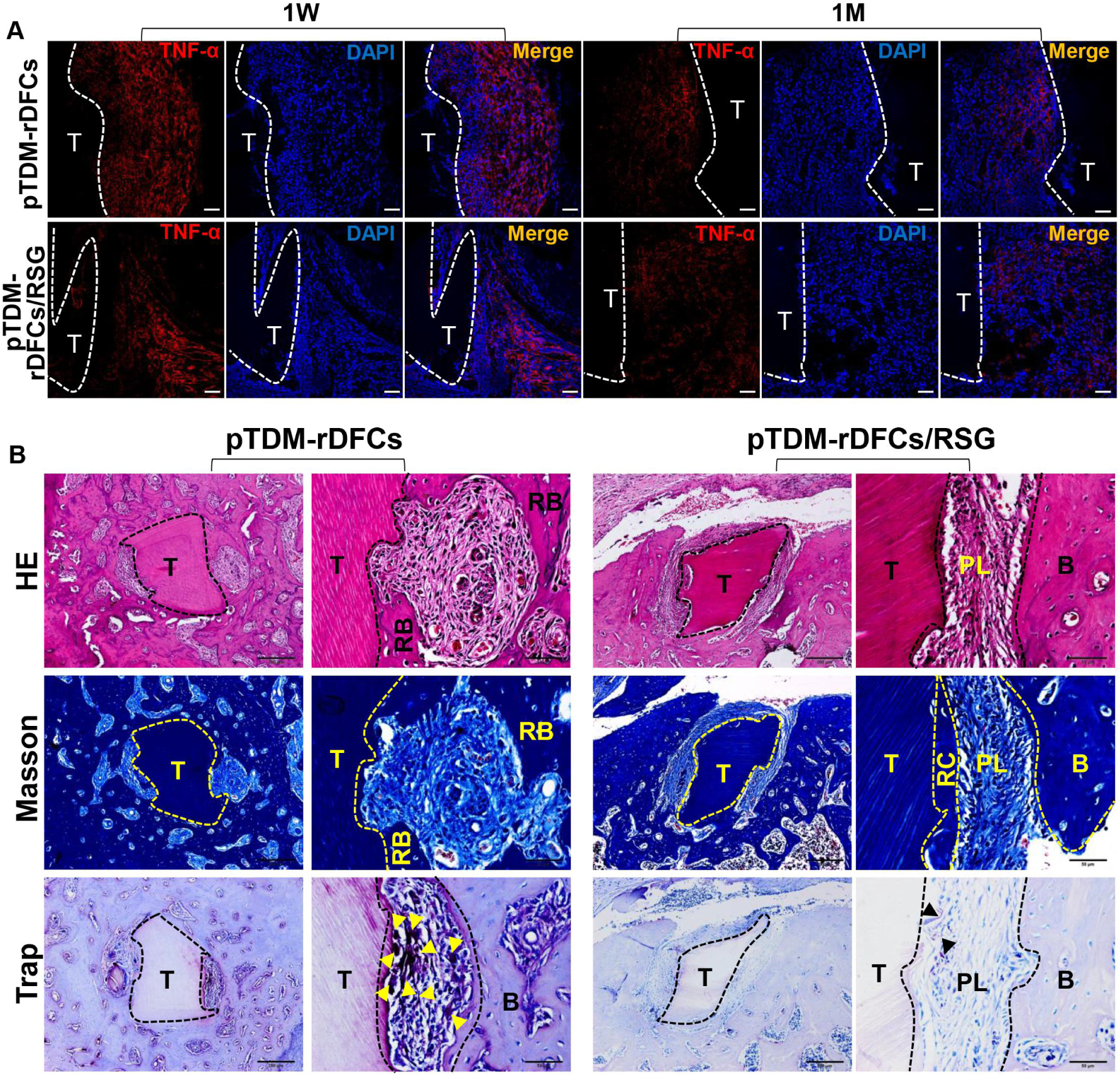
RSG inhibited osteoclasts differentiation while promoted PDL-like tissue regeneration in xenogeneic bioroots remodeling process by inhibiting inflammatory reaction. A, IF staining of TNF-α (red) around pTDM at 1 week and 1 month to detect inflammation level. T: pTDM. Scale bar = 100μm. B, H&E, Masson and trap staining of the xenogeneic bioroots in alveolar fossa of rats at 2 months post implantation. T: TDM; B: bone; RB: regenerated bone; RC: regenerated cementum; PL: periodontal-like tissues; OC (yellow and black arrows).

### RSG prevented xenogeneic bioroots resorption under an OS injury microenvironment

To further confirm the mechanism of the promotion effect of RSG on PDL-like tissue regeneration, xenogenic bioroots treated with or without RSG were subcutaneously implanted into the back of SOD1^-/ -^ C57 mice (***Figure 6A***). Two months after implantation, an obvious collagen matrix dissolution of pTDM accompanied by a low collagen fiber regeneration efficacy, a decreased periostin level and an elevated TNF-α expression were detected in xenogenic bioroots implanted in SOD1^-/ -^ C57 mice. On the contrary, when administrated with RSG, the transplanted pTDM maintained structural integrity without any collagen matrix dissolution or resorption. Other than that, large amounts of collagen fibers arranged perpendicular to pTDM were regenerated as shown in orthotopic implant model, demonstrating the importance of RSG in controlling ROS for enthesis regeneration in xenogenic bioroots (***Figure 6B and 6C***). Compared to the general view in SOD1- / - C57 mice with RSG treatment, the transplanted xenogenic bioroots without RSG treatment showed less collagen fiber encapsulation and less newly regenerated angiogenesis on the surface of pTDM (***Figure 6C***).

**Figure 6.**
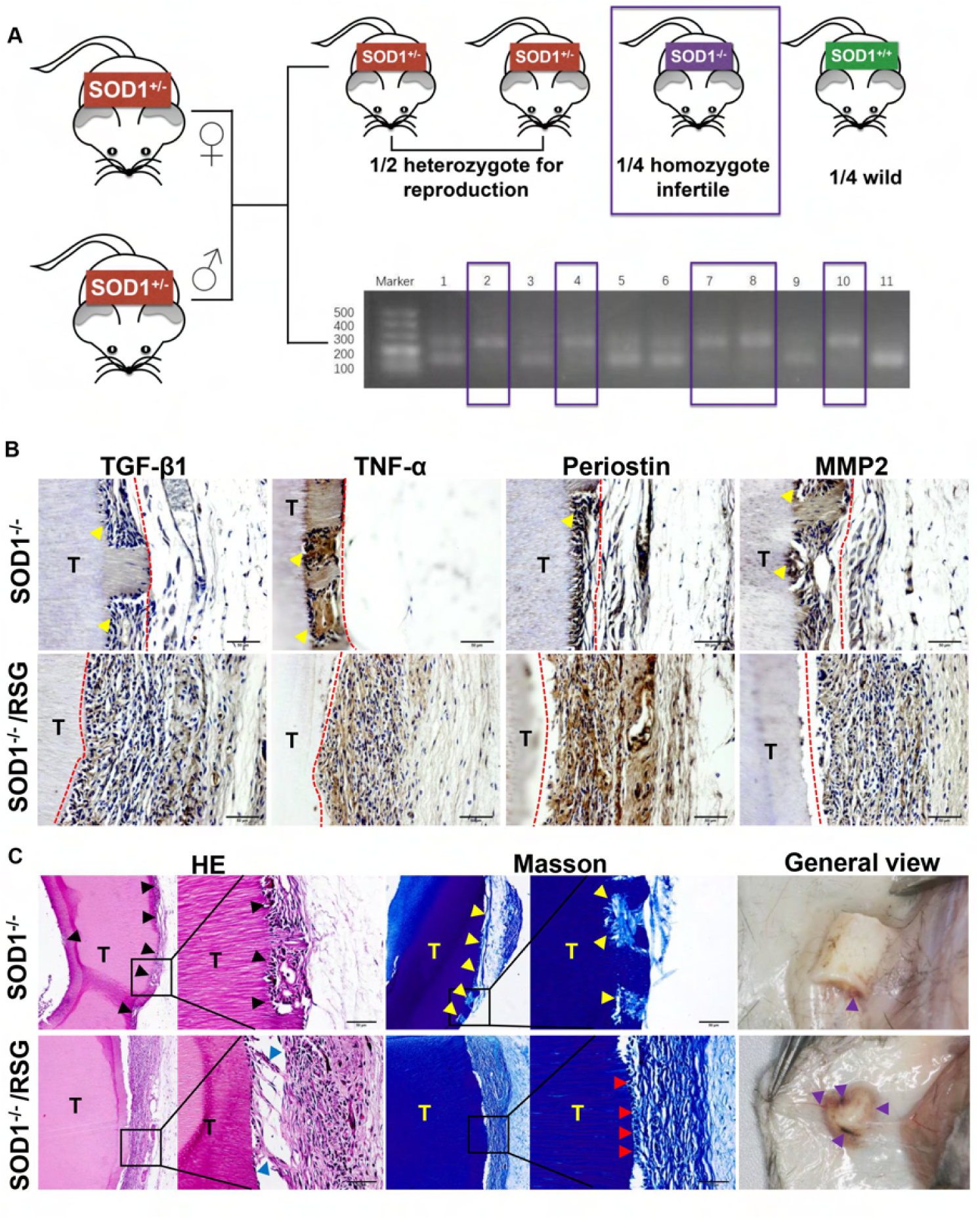
RSG prevented xenogeneic bioroots resorption under an OS injury microenvironment. A, clean homozygote SOD1 knockout mouse was defined by agarose gel electrophoresis. DNA marker (DL500) was made as the reference for DNA molecular weight; DNA bands between 100-200bp was defined as SOD1 wt genotype mice (sample 9 and 11), 200-300bp was defined as SOD1 KO genotype mice (sample 2,4,7,8 and10), 100-200bp and 200-300bp coexistence was defined as heterozygous mice (sample 1,3,5 and 6). B, IHC staining of the xenogeneic bioroots subcutaneously transplanted on the back of SOD1^-/-^ C57 mice. T: TDM. C, H&E, Masson staining and general view of the xenogeneic bioroots subcutaneously transplanted on the back of SOD1^-/-^ C57 mice. T: TDM; dissolved pTDM (black and yellow arrows); regenerated enthesis (red arrows); regenerated blood vessels (purple arrows).

## Discussion

Dental follicle stem cells are a kind of adult stem cells with multidirectional differentiation potential to become osteoblasts, adipoblasts and neuroblasts (***Yang et al., 2012***), which can be easily obtained from the third molar germs from adult subjects for the natural and normal development of teeth. Under an appropriate odontogenic induction environment, DFC is of great possibility to differentiate into periodontal ligament like tissues (***Yang et al., 2012***). pTDM, similar to the human derived TDM (hTDM), characterized by lax and porous structure on its surface (***Li et al., 2017***), will be the main source of biological scaffold to provide structural strength to transplant DFC for constructing xenogenic bioroots, considering its superiority in continuous release of the odontogenic differentiation related proteins like DMP-1/DSPP (***Li et al., 2011***). DMP-1/DSPP is considered to be an important protein in odontogenesis, which regulates cell mineralization (***Chen et al., 2020; Han et al., 2020***). When DMP-1/DSPP was knocked out, an obvious defect was observed on the surface of dental crown with dentin hypoplasia and tooth surface depression (***Saito et al., 2020***). Therefore, pTDM can be served as an ideal biological inducing material for guiding the odontogenic differentiation of DFC, suggesting the potential of constructing xenogenic bioroots by DFC wrapped with pTDM.

OS can cause significant oxidative damage on DNA, protein and lipid, directly causing tissue pathological changes. Therefore, ROS level can be evaluated by measuring the level of free radical metabolites and the antioxidant enzymes. 8-OHdG is a product of DNA oxidative damage and is often used as a marker for detecting ROS level and DNA mutation for early diseases diagnosis (***Xia et al., 2017***). Clinical studies have shown that 8-OHdG can be used as an important oxidative marker for evaluating the ROS level to confirm the effect of antioxidant therapy on colorectal cancer (***Xia et al., 2017***) or cardiovascular diseases (***Lai et al., 2013***), suggesting the probability of 8-OHdG in evaluating ROS expression. 3-NT is a kind of oxidation protein products originated from Tyrosine (Tyr) by adding a nitro group (-NO) with nitrating agents and is considered to be a marker of the formation of reactive nitrogens (RNs) in vivo. Nitrosative stress is closely related to oxidative stress. On the one hand, under oxidative stress, ROS can combine with RNs to form strong oxidant and nitrating agent to react with nucleic acids, proteins or other biological macromolecules. On the other hand, the oxidation product of nitrite-NO, also can damage intracellular bio-proteins, leading to the pathological changes (***Sinem et al., 2010***). At present, it has been reported that 3-NT can be used as an important marker of nitrosative stress to evaluate the therapeutic effect of antioxidants on asthma-chronic obstructive pulmonary disease overlap (***Kyogoku et al., 2019***). Therefore, in this study, 3-NT and 8-OHdG are both used to evaluate the ROS level.

PPARγ receptor exists in the nucleus of various cells and plays an important role in multiple physiological and pathological processes (***Rangwala et al., 2004; Rosen et al., 1999***). When activated, the microenvironment can be well regulated with an up-regulated expression of M2 macrophage polarization and a down-regulated level of inflammation, maintaining the immunity homeostasis (***Deng et al., 2019***). however, when PPARγ receptor was knocked out, the immunomodulation effect of macrophage was deprived with obvious proinflammatory factors production (***Silveira et al., 2019***). Therefore, thanks to its superiority in immune regulation, PPAR γ receptor agonist RSG has a wide application in xenotransplantation (***Lan et al., 2021; Han et al., 2020***). Consistent with this, in our study, RSG exhibited an anti-inflammatory effect on the H_2_O_2_ induced OS cells with decreased TNF-α expression but increased IL-4 concentration. Moreover, RSG also showed a positive role in scavenging ROS with increased cellular antioxidant capacity and a relative normal ultrastructure and mitochondrial membrane potential of mitochondria, suggesting the antioxidant capacity of RSG in the xenogenic bioroots system. In line with our study, previous research found that RSG could suppress the IRI during liver transplantation by promoting SOD1 expression (***Chen et al., 2017***). Besides, previous study also demonstrated that RSG showed an antioxidant effects on a various of chronic diseases associated with oxidative stress, such as Alzheimer’s syndrome, Parkinson’s syndrome and diabetes mellitus. Systemic assessment showed an obvious decrease in ROS concentration of these diseases in the serum after RSG treatment (***Kotha et al., 2021; Lee et al., 2019; Janani et al., 2015***), verifying the importance of RSG in inhibiting the excessive ROS production.

The features of the local immunological micro-environment largely determines the final regeneration results of the transplanted organs (***Jin et al., 2019***). Consistent redox imbalance can remarkably contribute to internal environment dysfunction along with exaggeration of inflammation, the leading cause of graft failure (***Kwiatkowska et al., 2021***). During renal replacement therapy, OS-induced damage often caused the development of chronic kidney disease (CKD)-related systemic complications with the transplanted renal fibrosis even death (***Colombo et al., 2020***). However, when the OS was suppressed with decreased ROS concentration by ozone, damages to renal were obviously slighter, promoting the transplanted renal survival (***Wang et al., 2018***). Similarly, in the heart transplantation, OS-derived damage contributes to progressive endothelial damage in the coronary arteries, leading to cardiac allograft vasculopathy initiation and progression (***Kargin et al., 2018***), greatly affecting the tissue remodeling process and function (**Schimke et al., 2000**). Recently, our study verified an increased regeneration efficiency of the allogenic bioroots with ROS controlling through administrating with NAC (***Zhang et al., 2021***). Therefore, antioxidant therapy should be a significant component of the patients with organ transplantation (***Szczurek et al., 2020***). Consistent with these studies, in this study, we also found an obvious up-regulated concentration of ROS after xenogenic bioroots implanted, resulting in a poor regeneration outcome accompanied by large amounts of bone adhesion and osteoclasts formation. However, when the intracellular ROS level and the inflammation expression in the transplantation microenvironment were well controlled by RSG, the organ survival rate was promoted significantly.

SOD1, as a subtype of the superoxide dismutases (SODs) with a surprisingly high cellular concentration (***Banci et al., 2013***), can preserve the ROS concentration at a normal level together with the other two antioxidant enzymes GSH / GPX by forming an antioxidant enzyme system. Previous researches found that SOD1 knockout mice exhibited excessive ROS accumulation and defects on multiple system, resulting in dysplasia or sickness (***Pouyet et al., 2010; Jaarsma et al., 2001; Hashizume et al., 2008; Hang et al., 2013***). Recently, our previous study have just found an obvious ROS aggregation in SOD1^-/-^ mice model with significant effects on mandible development. With the SOD1^-/-^ mice growing up, ROS concentration peaked at 6 months with an obvious up-regulated H_2_O_2_ expression and decreased bone formation rate (***Zhang et al., 2017***), reminding us that 6 months years old SOD1^-/-^ mice can be optimal for contributing OS model, with a potential to affect the tissue development and regeneration process. Therefore, for further detecting the antioxidant effect of RSG on xenogenic bioroots regeneration process, subcutaneous transplantation model was established in 4-month-old SOD1^-/-^ mice and the observation time was extended to 2 months. Consistent with the results got in orthotopic transplantation model of SD rats and the findings in other organs transplantation (***Elshazly et al., 2019***), RSG treatment was effective in promoting organ regeneration under OS microenvironment with less pDM collagen decomposition and inflammatory factors secretion but more periostin expression, verifying the antioxidant role of RSG.

## Conclusions

PPARγ agonist RSG can significantly alleviate the transplanted xenografts-derived OS damage by maintaining intracellular/immunity homeostasis, greatly promoting the regeneration outcomes and preserving the organs’ function.

## Materials and Methods

### Study approval

All animal experiments and procedures in this study were in accordance with institutional guidelines on animal welfare and were approved by the Institutional Laboratory Animal Care and Use Committee of Sichuan University (permit no. WCHSIRB-D-2019-063).

### pTDM preparation

Xenogeneic teeth, harvested from the freshly extracted deciduous incisor teeth of porcine, were made into pTDM (porcine derived treated dentin matrix) following gradient demineralization by 17%, 10% and 5% ethylene diamine tetraacetic acid (EDTA, Sigma-Aldrich, St. Louis, MO, USA) as previously reported (***Lan et al., 2021***). For pNDM (porcine derived natural dentin matrix), no demineralization was proceeded. All the obtained xECM were prepared for later experiment after performing ethylene oxide disinfection following freeze-dried for 8 hours.

### Biophysical properties of pTDM

SEM was used to determine the surface morphology of pTDM or pNDM at 15 kV. pTDM extracts were collected at different times after culturing in the saline with the weight/ volume ratio of pTDM to solvent 1:5 (g : ml) for protein release detection. At each time-point, equal volume of fresh saline were added into the dishes after the extracts were totally taken away.

### Cell culture and identification

rDFC (rat derived DFC) was isolated from the mandibular 1^st^ molar of 3-5 days old SD rats and cultured in a complete α-MEM medium containing 10% FBS (HyClone) and 1% of 100 U of penicillin and 100 μg/mL streptomycin at 37°C with 5% CO_2_ incubator. rDFC of 2-5 passage were seeded on 12-well plates at the number of 1×10^4^ for following experiments. For rDFC sheets formation, 50 mg/ml ascorbic acid was extra added into the medium to promote more extracellular matrix secretion when rDFC reached 70% confluence. For osteogenesis, 5 mM L-glycerophosphate (Sigma-Aldrich), 100 nM dexamethasone (Sigma-Aldrich), and 50 mg/ml ascorbic acid were supplemented into the medium for 21 days’ culturing. Subsequently, alizarin red S (ARS) staining was carried out after the cells were fixed with 4% paraformaldehyde (PFA). For neurogenesis, 2%DMSO (Amresco), 200μM Butylated hydroxyanisole (Sigma-Aldrich), 25 mM KCl(bdhg.company, China), 2 mM Valporic acid sodium salt (Sigma-Aldrich), 10mM forskolin (Sigma-Aldrich), 1mM hydrocortisone(aladdin), and 5μg/ml insulin (Novo Nordisk) were additionally imparted for culturing 2 hours. Afterwards, IF staining with βIII Tubulin (abcam, ab78078) was performed. Briefly, cells were fixed with 4% PFA at room temperature for 15 minutes after washing with PBS for three times, then permeabilized with 0.25% Triton-X and blocked with 5% BSA. Primary antibody against βIII Tubulin at a dilution of 1:100 was added onto the fixed cells and incubated overnight at 4 °C. DAPI was used to stain nuclei finally. All the IF staining in this study were all in accordance with this procedure (primary antibodies: anti-Vimentin (santa cruz, sc-6260), anti-CK14 (millipore, MAB3232), anti-Alex Fluor 555 Phalloidin (CST,#8953)).

### Cell proliferation, grouping and migration

CCK-8 test was carried out to assess the cells’ proliferation. Different concentrations of H_2_O_2_ and RSG were separately added into the 96-well plates to culture the cells with the counting number of 4000 per well initially. To detect the protection effect of RSG on the H_2_O_2_-induced damaged cells, rDFC was pretreated with 10uM RSG for 4 h before 25uM H_2_O_2_ was supplemented into the medium. With another 24 h culture, 10% (v/v) CCK-8 medium was added into each wells to culture the cells for 1.5 h at 37 °C for detecting the cell viability under different induced conditions. Cell viability was calculated based on the optical density (OD) value measured at 450nm by absorbance microplate reader following the formula: (OD value in control -OD value in blank) / (OD value in experiment - OD value in blank). Cell migration was assessed through scratch test when the cells were fusion with different chemical regents stimulation. For cell grouping, rDFC in rDFC/H_2_O_2_ group was treated with 25uM H_2_O_2_ for 24 hours while cells in rDFC/RSG/H_2_O_2_ group was pre-treated with 10uM RSG for 4 hours before cultured with 25uM H_2_O_2_. Pipette tip was used to make a scrape on the monolayer cell and the non-adherent cells were removed from the plates with PBS washed for three times. The cell morphology was recorded under light microscopy (Nikon, Japan) at 0 h, 24 h, and 48 h for cell proliferation and migration evaluation.

### ARS staining and RT-PCR

For evaluating the mineral nodule formation, ARS staining was performed after the cells were fixed with 4% PFA with culturing in the osteogenic medium for 21 days. After being cultured in the osteogenic medium for 7 days, rDFC was collected for assessing the expressions of the osteogenic and odontogenic differentiation-related genes or proteins as previously described (***Lan et al., 2021***). The specific primer information was shown as follows (5’-3’): GAPDH (Forward: TATGACTCTACCCACGGCAAG; Reverse: TACTCAGCACCAGCATCACC); COL-1 (Forward: TGCTGCCTTTTCTGTTCCTT; Reverse: AAGGTGCTGGGTAGGGAAGT); Runx2 (Forward: TCATTTGCACTGGGTCACAT; Reverse: TCTCAGCCATGTTTGTGCTC); ALP (Forward:CGTTGACTGTGGTTACTGCTGA; Reverse: CTTCTTGTCCGTGTCGCTCAC); Periostin (Forward:TTCGTTCGTGGCAGCACCTTC; Reverse: TCGCCTTCAATGTGGATCTTCGTC).

### Intracelluar ROS detection

For detecting intracelluar ROS level and inflammation expression, 2 ‘, 7’ - dichlorofluorescein diacetate (DCFH-DA, Sigma, America) staining, JC-1 (Beyotime, China) staining, transmission electron microscope (TEM) test and Elisa assay were conducted as previously described (***Lan et al., 2021***). In brief, cells were stained with JC-I or 10uM DCFH-DA in dark at 37 °C for 20 minutes for confocal microscope observation. For TEM detection, cells were fixed with 0.5% glutaraldehyde, dehydrated in gradient alcohol and permeated and embedded with resin. Consecutive 60-um-thick horizontal sections were obtained and double stained with uranium acetate and lead nitrate for 15 min for ultrastructural observation of mitochondria. Cell supernatants were collected for elisa test.

### Animal models and surgery

pTDM/rDFCs group and pTDM-RSG/rDFCs group were included in animal experiments. pTDM was cultured in 200 uM RSG solution at 37°C with 5% CO_2_ for 2 days and wrapped with the 5 uM RSG pretreated rDFCs, composed of pTDM-RSG/rDFCs group.

### Transplant the xenogeneic bioroots into an orthotopic implant model of SD rats

pTDM was prepared into a uniform size of 1 mm × 1 mm × 2 mm for orthotopic implant after ethylene oxide disinfection. A total of 48 Sprague-Dawley rats (8 weeks old, females, 230-260 g), purchased from Chengdu Dossy Biological Technology Company, were performed surgery. After anesthetization with 2% pentobarbital sodium (3 ml.kg − 1) and Zoletil 50 (50 mg.kg−1), skin, muscles and mucous membrane were blunt separated from the corner of the mouth until the bilateral 1^st^ molars could be directly seen. The xenogeneic bioroots, pretreated with or without RSG, were transplanted into the mesiobuccal roots of the freshly extracted 1^st^ molars. Then, the buccal and palatal mucosa were sutured to ensure that the implanted site could be completely covered. The muscle and skin in the open wounds were subsequently sutured in layers with 6-0 silk. At last, rats were injected with penicillin (1 U/200 g) and fasted for 10 hours postsurgery. 3-5 rats in each group were randomly selected and sacrificed at 1 week, 1 month and 2 months post implantation. Maxilla, including the first molar extraction socket, was harvested and fixed in 4% PFA at 4°overnight, and then demineralized with 10% EDTA for 2 months. Paraffin sections and frozen sections were prepared for subsequent HE, Masson, trap staining and IF staining as previously described (***Lan et al., 2021***). Briefly, frozen sections were washed by PBS for 3 times, repaired with antigen repair solution for 10 minutes, penetrated with 0.25% triton-100 for 10 minutes, blocked with 5% BSA for 1 hour, separately incubated with the primary antibody anti-4 Hydroxynonenal (abcam, ab48506) and anti-TNF α (santa cruz, sc-52746) at a dilution of 1:200 overnight at 4°C, subsequently cultured with the corresponding second fluorescent antibody for 1 hour, and finally stained with DAPI.

### Tissue homogenates extraction

This experiment was carried out as previously described (***Lan et al., 2021***). Briefly, SD rats were sacrificed at 7 days post surgery under general anesthesia and subsequently, maxilla was separated; then, the transplantation along with the surrounding fibrous tissues were obtained for further experiments. Precooled normal saline was used to rinse the samples for removing the red blood cells as much as possible. After, the 1:100 diluted PMSF was mixed with the tissue at the volume ratio of 9:1 to grind into tissue homogenate. Supernatants of the specimens were collected by centrifugation at 12000 rpm for 15 minutes at 4°C for subsequent Elisa analysis (8-OHdG/3-NT Elisa kit, Deco, Cat. #3455/ 3305).

### Transplant the xenogeneic bioroots into a subcutaneous implant model of SOD1^-/-^C57 mice

pTDM was shaped into a hollow circular column (3 mm × 2 mm × 4 mm) for subcutaneous implant after ethylene oxide disinfection. Shanghai Model Organisms Center, Inc. was entrusted to establish an animal disease model of B6;129S-Sod1^tm1Leb^J (SOD1^-/-^) of C57 mice. After anesthetized with isoflurane, the prepared xenogeneic bioroots with or without RSG pretreated were implanted subcutaneously on the back of 4-month-old SOD1^-/-^ C57 mice. The mice were killed 2 months after implantation and the samples were fixed at 4°C in 4% PFA overnight, and then demineralized with 10% EDTA for 4 months. Paraffin sections were prepared for subsequent histological staining as previously described (***Lan et al., 2021***). Anti-TGFβ1 (abcam, ab92486), anti-Periostin (H-300) (santa cruz, sc-67233) and anti-MMP2 (abcam, ab92536) primary antibodies were applied at a dilution of 1:200.

### Statistical analysis

All data are presented as the mean ± standard deviation (M ± SD). The Student’s t-test was used to assess differences between two groups. One-way analysis of variance (ANOVA) was performed to evaluate the discrepancies among multiple groups. All in vitro or in vivo work was represented at least three biological replicates by independent experiments.

## Acknowledgments

This work was supported by the Nature Science Foundation of China (31971281), Innovative Talents Program of Sichuan Province (2022JDRC0043), Research and Develop Program, West China Hospital of Stomatology Sichuan University (RD-03-202106)

## Author contributions

(I) Conception and design: Tingting Lan; (II) Administrative support: Weihua Guo; (III) Provision of study materials: Jie Chen, Xiaoli Yin; (IV) Collection and assembly of data: Xue Han, Yuchan Xu and Xiaoli Yin; (V) Data analysis and interpretation: Fei Bi, Tingting Lan and Weihua Guo; (VI) Manuscript writing: All authors; (VII) Final approval of manuscript: All authors.

